# Immune Responses Targeting Homocitrullinated Peptides Drive Rheumatoid Arthritis-like Pathology in HLA-DR4 Transgenic Mice

**DOI:** 10.1101/2025.09.17.676523

**Authors:** Gabrielle Buckley, Sheri Saunders, Ewa Cairns, Lillian Barra

## Abstract

Rheumatoid arthritis (RA) is a systemic autoimmune disease characterized by persistent synovial inflammation and progressive joint destruction. A hallmark feature of RA is the presence of anti-citrullinated protein autoantibodies (ACPAs), which have been extensively studied in disease pathogenesis. The development of these autoantibodies is strongly associated with the expression of HLA-DR4, containing the shared epitope. More recently, anti-homocitrullinated protein autoantibodies (AHCPAs) have also been implicated in RA and demonstrate a similar genetic association. Therefore, this study aimed to evaluate the role of homocitrulline immune responses in RA pathogenesis. HLA-DR4 transgenic (DR4tg) and B6 mice were immunized with a synthetic homocitrullinated peptide (HomoCitJED) or PBS followed by a booster 21 days later. Beginning on day 75 post-primary immunization mice received knee intra-articular (i.a.) injections of HomoCitJED once a week for three consecutive weeks. Swelling post i.a. injection was measured using digital calipers. IgG antibodies against homocitrullinated peptides/proteins were measured using enzyme-linked immunosorbent assay (ELISA). Knees were sectioned and analyzed for histopathological damage and citrullinated/homocitrullinated protein levels. HomoCitJED immunized DR4tg mice exhibited significantly greater swelling and incidence of anti-HomoCitJED and -homocitrullinated fibrinogen IgG autoantibodies. Additionally, 30% had responses against CitJED, a synthetic citrullinated peptide. HomoCitJED immunized DR4tg mice also experienced greater total histopathological damage; however, the immunizations/injections did not affect citrullinated/homocitrullinated protein levels. In conclusion, homocitrulline-specific immune responses can induce RA-like disease in DR4tg mice, supporting the pathogenic role of AHCPAs in RA.

## 1. Introduction

Rheumatoid arthritis (RA) is a chronic, systemic autoimmune disease characterized by persistent inflammation of the synovial joints. Affecting about 1% of the global population, RA ranks among the most common autoimmune diseases [1,2]. Its onset usually occurs between the ages of 30 and 50, with women three times more likely to develop the disease [3,4]. RA has a predilection for the small peripheral joints of the hands and feet but can also involve the larger joints. If untreated, RA can result in permanent joint damage and functional impairment [5]. Disease progression is driven by persistent synovial inflammation that leads to synovial infiltration of joint tissues (pannus formation) with cartilage and bone damage [1,6]. The underlying inflammatory processes are influenced by genetic predisposition, which accounts for nearly 60% of the risk of developing the disease, while environmental and lifestyle factors also play critical roles [7]. The greatest genetic risk factor for RA development is the presence of major histocompatibility complex (MHC) class II molecules encoded by HLA-DRB1 alleles containing the shared epitope (SE), a conserved five-amino-acid sequence motif located within the peptide-binding groove of the DRB1, including HLA-DRB1*04 (HLA-DR4) and HLA-DRB1*01 (HLA-DR1) [8,9].

While the exact cause of RA remains unknown, its onset is thought to involve a breakdown in immune tolerance to post-translationally modified proteins, often citrullinated and homocitrullinated proteins (CitP/HomoCitP) that are present in the joint. Citrullination is a post-translational modification of arginine, catalyzed by peptidylarginine deiminase (PAD) enzymes, while homocitrullination is a modification of lysine, occurring through the action of myeloperoxidase (MPO) or cyanate-dependent chemical processes [10]. This breakdown in tolerance leads to the formation of anti-citrullinated protein autoantibodies (ACPAs) and anti-homocitrullinated protein autoantibodies (AHCPAs) [10]. The development of ACPA is closely associated with the SE through altered antigen presentation. Hill *et al.* previously showed that HLA-DR4 and HLA-DR1 had a higher binding affinity for a vimentin-derived peptide when it was citrullinated compared to the native peptide, but not in HLA-DRB1 alleles lacking the SE [9]. Based on *in silico* modelling, a similar increased binding affinity is thought to occur for HomoCitP, which differs from citrulline by a single carbon atom [11].

ACPAs are highly specific to RA and appear to be involved in the disease, as injection of isolated human ACPAs into mice induces pain, osteoclast activation, and bone erosion, resembling the disease manifestations observed in humans [12–15]. The role of AHCPAs in RA pathogenesis is less studied; however, they are specific to RA and Scinocca *et al.* found anti-homocitrullinated fibrinogen autoantibodies in ∼50% of RA patients [11]. Moreover, Shi *et al.* found that 16% of RA patients who were ACPA negative were AHCPA positive [16]. Recently, Lac *et al.* demonstrated that HLA-DR4-transgenic (DR4tg) mice immunized with a HomoCitP developed AHCPAs, providing a clinically relevant animal model for studying their arthritogenic properties [17]. Therefore, our study aimed to evaluate the ability of immune responses against homocitrulline to induce arthritis in DR4tg mice by targeting them to the joint via intra-articular (i.a.) injections of a HomoCitP.

## 2. Materials and methods

### 2.1 Peptide and Protein Antigens

The antigens used in this study include: 1) citrullinated human fibrinogen (CitFib), 2) homocitrullinated human fibrinogen (HomoCitFib), 3) citrullinated JED (CitJED), 4) homocitrullinated JED (HomoCitJED), and 5) unmodified human fibrinogen (VWR). Citrullination, with PAD in 0.1 M Tris-HCL (pH 7.4), 10 mM CaCl_2_, and homocitrullination, with 0.1 m potassium cyanate in 0.15 m sodium phosphate buffer of fibrinogen were performed in house as previously described by Hill *et al.* [18] and Scinocca *et al.* [11]. CitJED and HomoCitJED peptides were synthesized by Creative Peptides (Shirley, NY, USA) and contain 18 amino acid residues, 9 of which are citrulline or homocitrulline, respectively. The remaining residues that comprise the backbone are identical between peptides [11,17,19].

### 2.2 Mice

All experiments performed in this study adhered to the Canadian Council on Animal Care guidelines and were approved by the Animal Care and Use Committee at Western University (AUP 2022-150). Male and female DR4-IE transgenic, murine MHC class II-deficient (DR4tg) mice (bred in-house) [20,21] and C57Bl/6 (B6) mice (Jackson Laboratory, Maine, USA) were housed in a pathogen-free barrier facility. Mice 8-12 weeks old were subcutaneously immunized in the flank with 100 µg HomoCitJED (dissolved in PBS), emulsified in 2.0 mg/mL complete Freund’s adjuvant (CFA) (*Mycobacterium tuberculosis* H37RA (Fisher Scientific)) [17,21]. Control mice were immunized with PBS in CFA (2.0 mg/mL). Booster immunizations were given 21 days later in incomplete Freund’s adjuvant (IFA) [17,21]. To induce arthritis, mice received on day 75 an i.a. injection of 25 ug HomoCitJED (dissolved in PBS) in the left knee and PBS in the right knee. To control for any effects from the i.a. injections, some PBS immunized mice received i.a. injections in the left knee and no injection in the right knee. Subsequent i.a. injections occurred at one-week intervals for a total of three injections. Prior to the start of this i.a. regimen saphenous blood was collected, which was continued at two-week intervals until the day of sacrifice. Day 75 was chosen for the beginning of the i.a. injection regimen as Lac *et al.* previously showed it to be when antibody responses peak and there was no difference in antibody levels [17] or histopathology at alternative timepoints (Supplementary Figure 1). Digital calipers were used to monitor knee swelling for at least three consecutive days following each i.a. injection, then once weekly following regimen completion. Swelling was considered to be a change in knee width > 0.2 mm, two standard deviations above the mean of naïve mouse knee measurements. Mice were sacrificed 137 days post-primary immunization.

### 2.3 Antibody Assays

Sera collected at various time points was screened for the presence of IgG antibodies towards HomoCitJED, CitJED, HomoCitFib, CitFib, and unmodified fibrinogen via enzyme linked immunosorbent assay (ELISA), as previously described by Lac *et al*. [17]. The sera were tested in duplicate with a coefficient of variation <20%. The positive cut-off optical density (OD) was 0.1, the ELISA’s lower detection limit (corresponding to 20.32 RU/mL, 2.86 RU/mL, and 12.05RU/mL for HomoCitJED, HomoCitFib, and CitJED, respectively) [4]. Antibody concentrations in RU/mL were determined through creation of a standard curve using a pooled positive reference serum.

### 2.4 Histopathological Analysis

Hind limbs were collected at sacrifice and fixed in 10% buffered formalin (Fisher Scientific) for 24 hours. The fixed tissue was then decalcified in 14% EDTA (Thermo Scientific) for two weeks at 4ᵒC, followed by dehydration. The knees were then embedded in paraffin and sagittally sectioned (5 µm). Serial knee sections were stained with hematoxylin and eosin (H&E) and toluidine blue. Blinded histopathological scoring for synovial inflammation, bone erosion, proteoglycan loss, and cartilage erosion was then performed following the SMASH protocol for Tg197 mice [6].

### 2.5 Immunofluorescent Staining

Serial tissue sections were permeabilized with 0.25% Triton X-100 (Sigma), followed by enzymatic antigen retrieval with trypsin (Abcam). Tissues were then blocked with 10% goat serum (Thermo-Fisher) prior to the application of a Rabbit IgG Polyclonal Isotype Control Antibody (Abcam) and Rabbit Anti-Carbamyl-Lysine (CBL) Polyclonal IgG Antibody (Cell Biolabs). Slides were incubated at 4ᵒC for 24 hours and then washed. Subsequently, a Mouse IgMκ Monoclonal Isotype Control Antibody (Abcam) and Peptidyl-Citrulline, Clone F-95, Monoclonal IgMκ Antibody (EMD Millipore) were applied and incubated at 4ᵒC for 24 hours. Next, Alexa Fluor® AffiniPureTM F(ab’)2 Fragment Goat Anti-Mouse IgM, µ chain specific, Polyclonal Secondary Antibody (Jackson) and Alexa Fluor® 647 AffiniPureTM F(ab’)2 Fragment Goat Anti-Rabbit IgG, Fc Fragment specific Secondary Antibody (Jackson) were applied for one hour. TrueView autofluorescence quencher and DAPI mounting media (Vector) were then applied.

Slides were imaged on the Nikon Laser Scanning Confocal Hybrid microscope with the following settings: DAPI (Power = 5, Gain = 74), FITC (Power = 2, Gain = 34), CY5 (Power = 2, Gain = 97). Regions of interest (ROIs) for bone marrow, cartilage, and synovium were defined using ImageJ software. Corrected Total Fluorescence was calculated using the following equation: Integrated Density of F-95/CBL – (Area x Mean Fluorescence of Isotype Control).

### 2.6 Statistics

Statistical analyses included the Kruskal-Wallis test with Dunn’s multiple comparison test for non-parametric data, and Welch’s ANOVA with Dunnett’s T3 multiple comparison test, ordinary one-way ANOVA with Dunnett’s multiple comparison test, or paired or unpaired t tests for parametric data. Welch’s ANOVA was applied for parametric data with a significant Barlett’s test for unequal variance, while ordinary one-way ANOVA was used for parametric data with a non-significant Barlett’s test. Contingency analyses used the Chi-square test. All analyses were performed in GraphPAD Prism 10; analyses were considered significant if p < 0.02, corrected for multiple comparisons.

## 3. Results

### 3.1 HomoCitJED immunized DR4tg mice develop anti-HomoCitJED and anti-HomoCitFib IgG autoantibodies

DR4tg and B6 mice were screened for IgG antibodies (Abs) against HomoCitP and CitP at various timepoints after immunization with HomoCitJED vs. PBS (Fig. 1). HomoCitJED-immunized DR4tg mice had significantly higher levels of anti-HomoCitJED IgG Abs over time than the PBS immunized DR4tg mice (p = 0.0002), but not HomoCitJED or PBS immunized B6 mice (p = 0.0821 and p = 0.0516, respectively) (Fig. 1A). 3/14 (93%) DR4tg mice immunized with HomoCitJED were found to be positive for anti-HomoCitJED IgG Abs, while only 3/6 (50%) and 0/4 (0%) HomoCitJED and PBS immunized B6 mice, respectively were considered positive (p = 0.0281 and p = 0.0003). Anti-HomoCitFib IgG Abs were assessed as a naturally occurring antigen. Significantly higher levels of anti-HomoCitFib IgG Abs were observed in HomoCitJED immunized DR4tg mice vs. PBS immunized DR4tg mice and HomoCitJED immunized B6 mice (p = 0.0041 and 0.0066, respectively). Furthermore, 13/14 (93%) HomoCitJED immunized DR4tg mice while 0/6 (0%) HomoCitJED and 1/4 PBS immunized B6 mice were considered positive for anti-HomoCitFib Abs, respectively (p < 0.0001 and p = 0.0040) (Fig 1B). Potential epitope spreading to citrullinated antigens was then assessed by looking at IgG antibodies towards CitJED: 4/14 (29%) and 1/6 (17%) HomoCitJED immunized DR4tg and B6 mice, respectively were considered positive (p = 0.5731) (Fig 1C). PBS immunized DR4tg and B6 mice were not found to be positive for IgG antibodies towards CitJED. Additionally, no IgG antibodies towards unmodified fibrinogen or CitFib were detected in DR4tg or B6 HomoCitJED immunized mice (Supplementary Figure 1). I.a. injection of HomoCitJED was also found to not alter anti-HomoCitJED IgG antibody levels, which were stable over time (Figure 1A).

**Fig 1.**
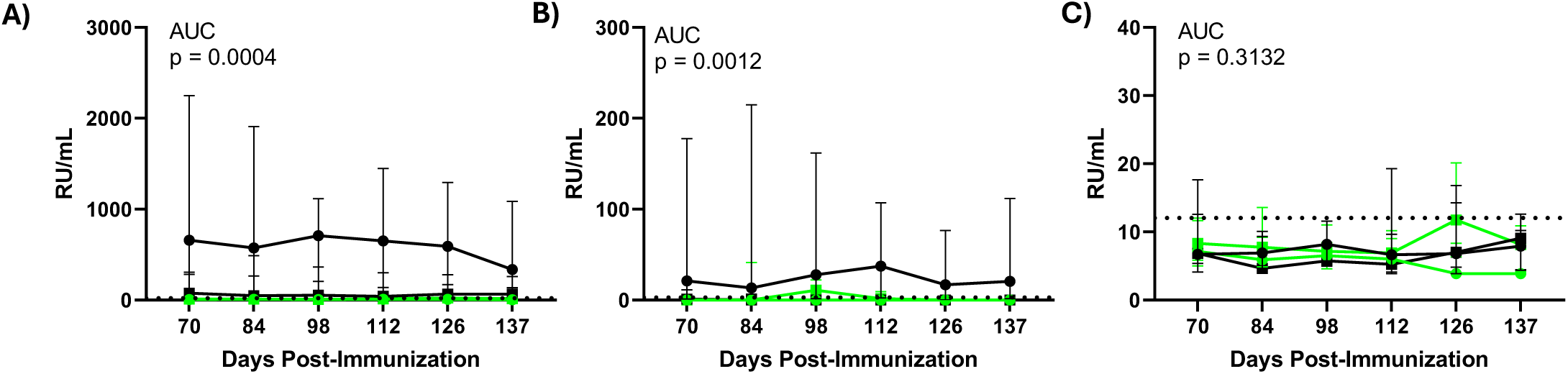
IgG antibody responses in DR4tg and B6 mice. Sera from DR4tg [HomoCitJED immunized (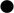), PBS immunized (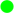)] and B6 [HomoCitJED immunized (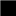), PBS immunized (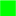)] mice were screened for (**A**) anti-HomoCitJED, (**B**) anti-HomoCitFib, and (**C**) anti-CitJED antibodies measured at the indicated time points post-immunization. All mice received i.a injections of HomoCitJED in the left knee and PBS in the right knee. Antibody levels, determined via ELISA, are shown in RU/mL. Each symbol represents the median value (IQR) with the positive cutoff shown by a black dashed line; N = 4-14. A Kruskal-Wallis statistical test of area under the curve was performed.

### 3.2 HomoCitJED immunized DR4tg mice develop knee swelling when given i.a. injections of HomoCitJED

DR4tg and B6 mice were assessed for changes in knee width post i.a. injection. HomoCitJED immunized DR4tg mice had significantly greater swelling after receiving HomoCitJED i.a. injections than PBS immunized DR4tg mice (p < 0.0001), HomoCitJED immunized B6 mice (p = 0.0076), and PBS immunized B6 mice (p <0.0001) (Fig. 2A). Furthermore, 14/14 (100%) HomoCitJED immunized DR4tg mice were considered significantly swollen compared to 3/6 (50%) HomoCitJED immunized B6 mice following HomoCitJED i.a. injections. PBS immunized mice did not develop significant swelling when given HomoCitJED i.a. injections. Additionally, no swelling was observed in the knee receiving PBS i.a. injections regardless of immunization (p = 0.3634; Fig 2B).

**Fig 2.**
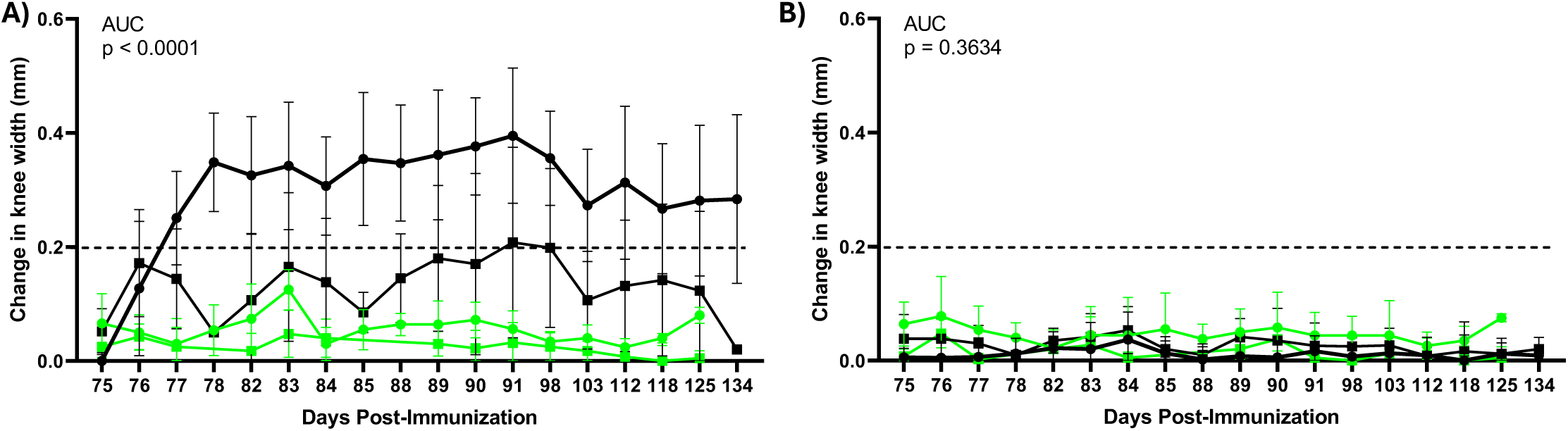
Joint swelling of DR4tg and B6 mice. Changes in knee width caliper measurements for the (**A**) right and (**B**) left knee were compared between DR4tg [HomoCitJED immunized (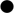), PBS immunized (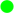)] and B6 [HomoCitJED immunized (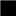), PBS immunized (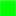)] mice at the indicated time points post-immunization. All mice received i.a injections of HomoCitJED in the left knee and PBS in the right knee. Each symbol represents the mean value (SD) with the swelling cutoff shown by a black dashed line; N = 4-14. A Welch ANOVA statistical test of area under the curve was performed.

### 3.3 HomoCitJED immunized DR4tg mice have histopathologic evidence of arthritis

Arthritis severity at sacrifice was determined through histopathological scoring of synovial inflammation, bone erosion, proteoglycan loss, and cartilage erosion. Mean (SD) of total histopathological scores, out of 12, for HomoCitJED immunized DR4tg, PBS immunized DR4tg, HomoCitJED immunized B6, and PBS immunized B6 were 2.5 (0.84), 1.2 (0.4), 1.4 (0.5), and 1.2 (0.4), respectively. The total histopathological scores were significantly more in HomoCitJED immunized DR4tg mice compared to PBS-immunized DR4tg mice and B6 mice (p = 0.0098, 0.0129, and 0.0084). The histopathological scores in the HomoCitJED immunized DR4tg mice were not found to be different between the right and left knees (p = 0.7675; Fig 3B); but male mice had greater severity than females (p = 0.0314; Fig 3C).

**Fig 3.**
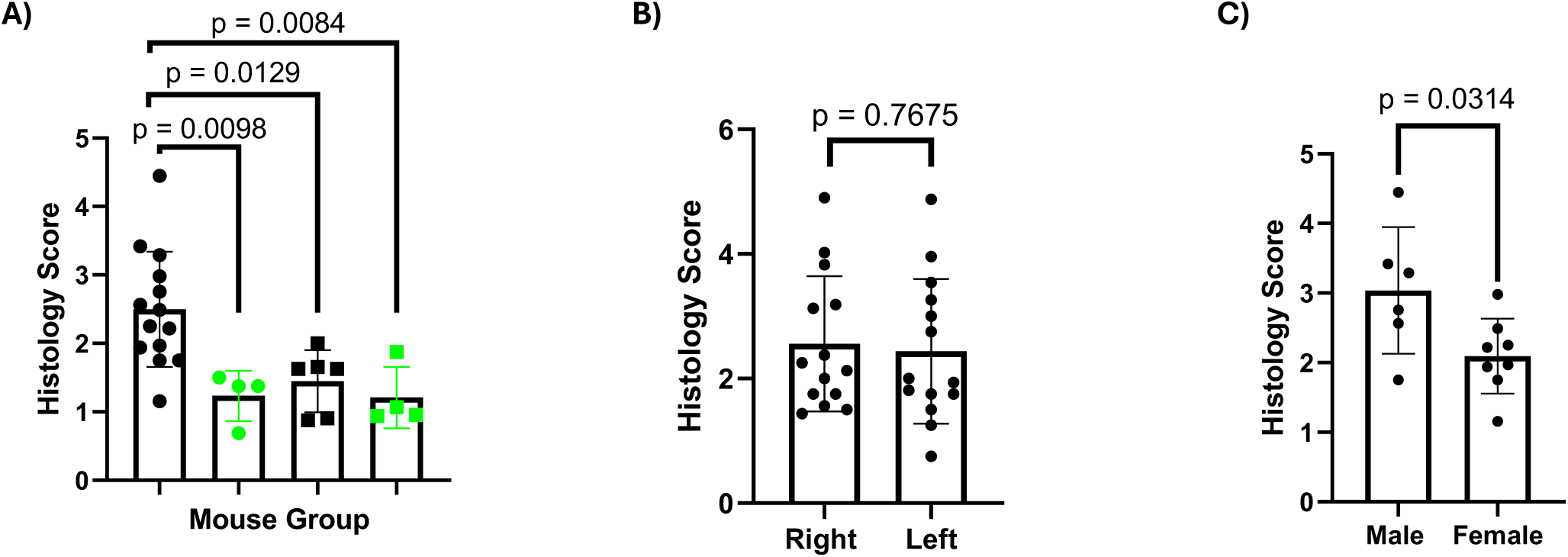
Histological analysis of arthritis induced by HomoCitJED in DR4tg and B6 mice. (**A**) Total histopathological scores was compared between DR4tg [HomoCitJED immunized (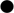), PBS immunized (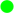)] and B6 [HomoCitJED immunized (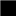), PBS immunized (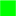)] mice. All mice received i.a injections of HomoCitJED in the left knee and PBS in the right knee. Histological scores from the right and left knee were averaged. Each symbol represents one mouse; N = 4-14. An ordinary one-way ANOVA statistical test was performed (p = 0.0015). (**B**) Comparison of total histopathological scores in the right and left knees of DR4tg mice immunized with HomoCitJED. Each symbol represents one knee; N = 14. A paired t-test was performed. (**C**) Comparison of total histopathological scores in male (N = 6) and female (N = 8) DR4tg mice immunized with HomoCitJED. Histological scores from the right and left knee were averaged, with each symbol representing one mouse. An unpaired t-test was performed. All graphs display the mean (SD).

Evaluation of the histopathological score sub-categories showed that synovial inflammation and proteoglycan loss were higher in DR4tg mice immunized with HomoCitJED, but did not reach statistical significance (Figure 4A-E and Figure 5A-E, respectively). Bone erosions were very minimal across all groups (Figure 4A-D, F). Similarly, cartilage erosion was found to be minor across all groups but trended to being higher in HomoCitJED immunized DR4tg mice (Figure 5A-D, F). The observed histopathological damage was not fully due to the i.a. injection regimen, as PBS immunized mice receiving i.a. injections of PBS in the left knee and no i.a. in the right were not significantly different (Supplementary Figure 3).

**Fig 4.**
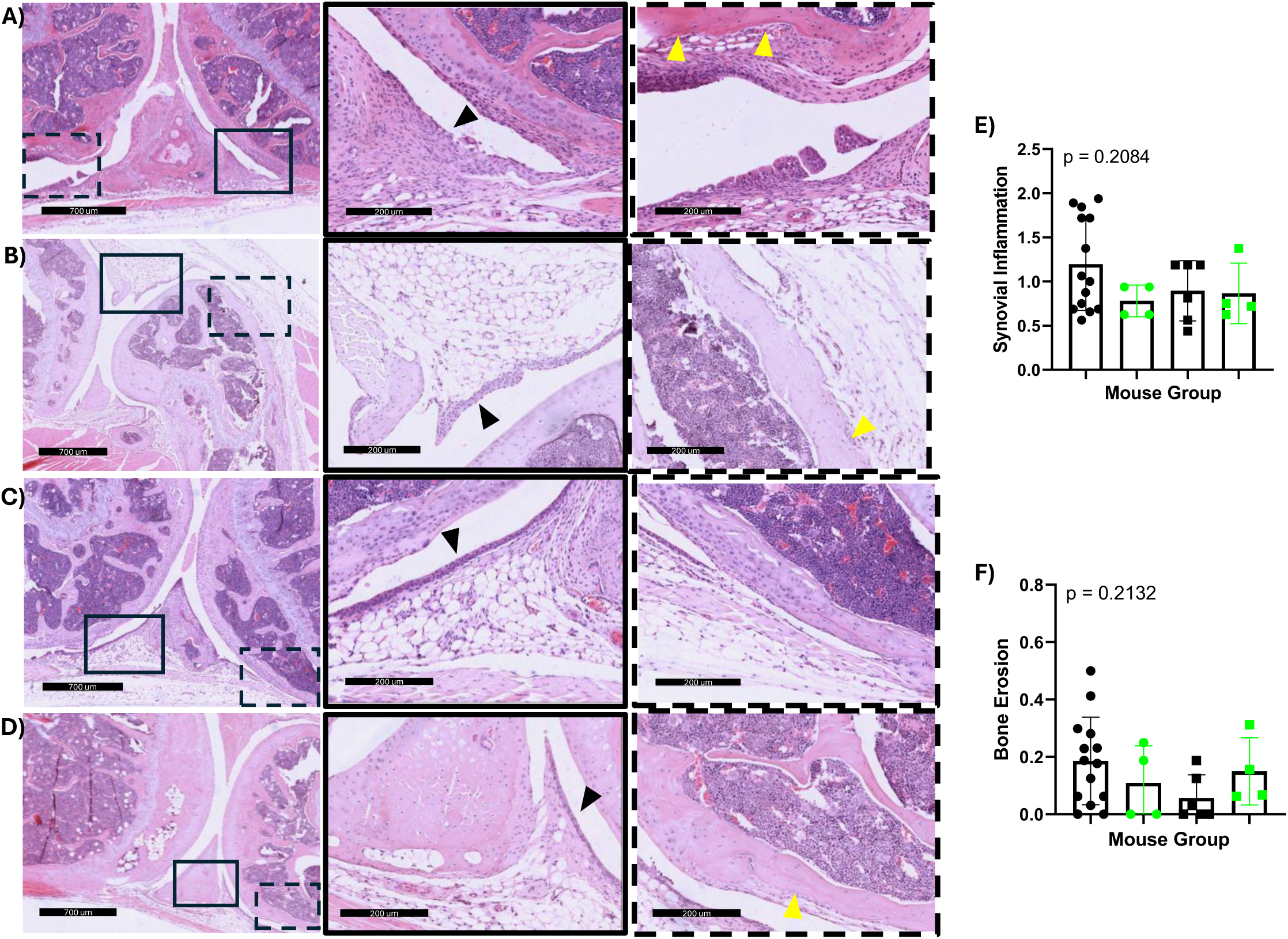
Histological analysis of synovial inflammation and bone erosion induced by HomoCitJED in DR4tg and B6 mice. Representative H&E images of (**A**) HomoCitJED immunized DR4tg mice, (**B**) PBS immunized DR4tg mice, (**C**) HomoCitJED immunized B6 mice, and (**D**) PBS immunized B6 mice. The images in the first column were obtained at 60x magnification (scale bar = 700 μm), with adjacent images enlarged at 240x magnification (scale bar = 200 μm). Histopathological features: synovial inflammation (black arrowhead); bone erosion (yellow arrowhead). (**E**) Synovial inflammation and (**F**) bone erosion histopathological scores were compared between DR4tg [HomoCitJED immunized (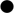), PBS immunized (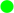)] and B6 [HomoCitJED immunized (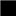), PBS immunized (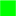)] mice. All mice received i.a. injections of HomoCitJED in the left knee and PBS in the right knee. Histological scores from the right and left knee were averaged. Graphs display the mean (SD), with each symbol representing one mouse; N = 4-14. A welch ANOVA statistical test was performed.

**Fig 5.**
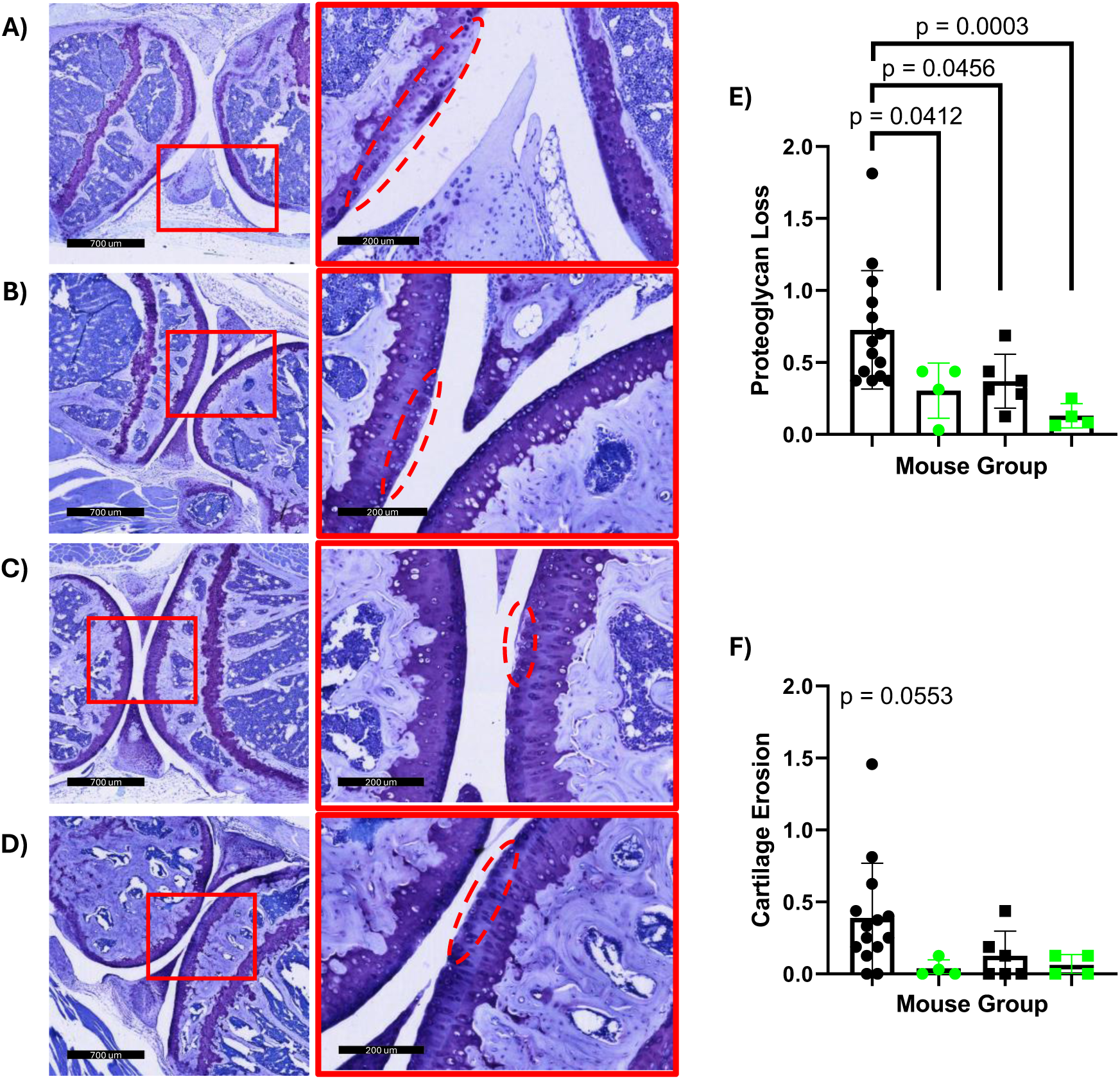
Histological analysis of proteoglycan loss and cartilage erosion induced by HomoCitJED in DR4tg and B6 mice. Representative toluidine blue images of (**A**) HomoCitJED immunized DR4tg mice, (**B**) PBS immunized DR4tg mice, (**C**) HomoCitJED immunized B6 mice, and (**D**) PBS immunized B6 mice. The images in the first column were obtained at 60x magnification (scale bar = 700 μm), with adjacent images enlarged at 240x magnification (scale bar = 200 μm). Histopathological features: proteoglycan loss (dashed red oval). (**E**) Proteoglycan loss and (**F**) cartilage erosion histopathological scores were compared between DR4tg [HomoCitJED immunized (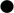), PBS immunized (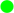)] and B6 [HomoCitJED immunized (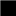), PBS immunized (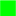)] mice. All mice received i.a. injections of HomoCitJED in the left knee and PBS in the right knee. Histological scores from the right and left knee were averaged. Graphs display the mean (SD), with each symbol representing one mouse; N = 4-14. A welch ANOVA statistical test was performed for proteoglycan loss (p = 0.0040) and cartilage erosion.

### 3.4 CitP and HomoCitP are present in the knees of DR4tg mice and are not affected by HomoCitJED immunization or i.a. injections

CitP and HomoCitP levels in the knees were determined through immunofluorescent staining. These modified proteins were observed in the joints of HomoCitJED immunized DR4tg mice, PBS immunized DR4tg mice, and naïve DR4tg mice (Figure 6A-C, Supplementary Figure 4A-C). Levels of CitP and HomoCitP were found not to be affected by HomoCitJED immunization or i.a. injection in the bone marrow, cartilage, or synovium (Supplementary Figure 5).

**Fig 6.**
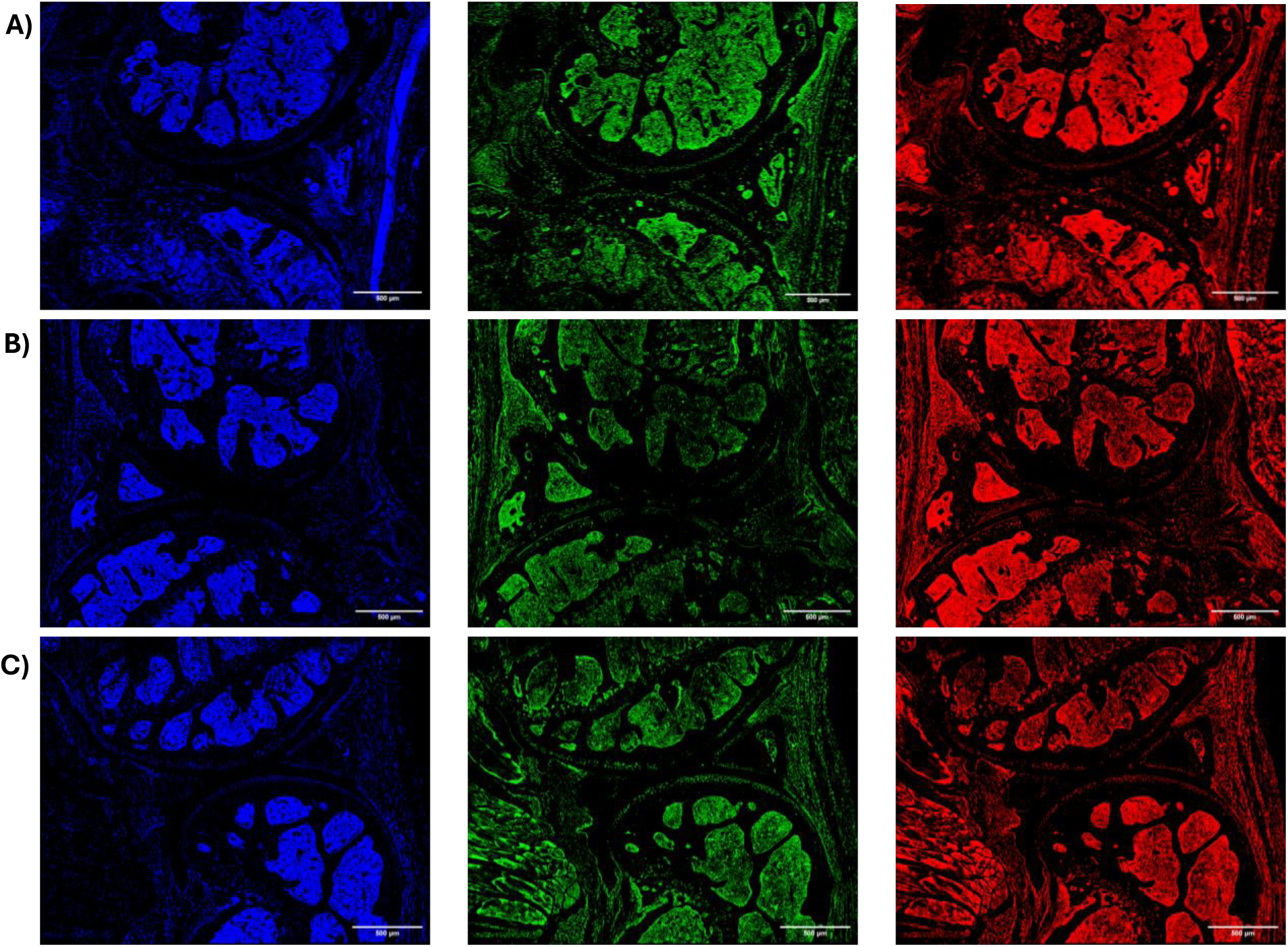
Representative immunofluorescent images of CitP and HCitP in joints of DR4tg mice. Representative images of (**A**) HomoCitJED immunized DR4tg mice, (**B**) PBS immunized DR4tg mice, and (**C**) naïve DR4tg mice. Joints were stained such that nuclei, CitP, and HCitP are DAPI (blue), FITC (green), and CY5 (red), respectively. Immunized mice received i.a. injections of HomoCitJED in the left knee and PBS in the right knee. The images are at 20x magnification; scale bar = 500 µm.

## 4. Discussion

Immune responses targeting HomoCitP have recently been implicated in the pathogenesis of RA [11]. CitP is a well-characterized autoantigen in RA that is the target of inflammatory immune responses associated with joint pain and bone erosions [13–15]. HomoCitP, which is structurally similar, may also contribute to similar processes, although its precise role remains to be fully established. In this study, we demonstrated that knee i.a. injections of a HomoCitP (HomoCitJED) following immunization with HomoCitJED in DR4tg mice is sufficient to induce joint pathology resembling the hallmark features of RA. These findings provide evidence for a mechanistic link between HomoCitP-directed immune responses and RA development in genetically susceptible individuals.

DR4tg mice immunized with HomoCitJED followed by i.a. injections of the peptide developed signs of inflammatory arthritis, including synovitis, bone erosions, proteoglycan loss, and cartilage erosion. Synovitis was the most prominent histopathological feature in our model, which has been shown to predict subsequent radiographic progression in RA [22]. In contrast, PBS immunized controls that received i.a. injections of HomoCitJED exhibited significantly lower arthritis histopathology scores and lacked detectable antibody responses, underscoring the importance of anti-HomoCitJED immune responses for disease induction. This is further supported by our observation that HomoCitP and CitP were detectable in the knees of DR4tg mice across all groups, including naïve animals, with no measurable differences in their levels. Consistent with this, high-performance liquid chromatography (HPLC) of human synovial tissue have demonstrated PAD expression in healthy joints [23]. MPO levels are generally low or undetectable in the synovial fluid [24]. However, homocitrullination in human skin has been shown to naturally increase over time, highlighting its physiological occurrence [25]. Together, these findings suggest that the presence of antigen alone does not result in RA-associated joint pathology.

Our findings are consistent with Mydel *et al.,* who observed that immunization with a synthetic filaggrin-based HomoCitP, followed by i.a. injection of its citrullinated form, induced erosive arthritis, whereas non-immunized mice receiving i.a. injections of the HomoCitP developed only mild synovitis with occasional bone erosions but no cartilage damage [26]. The histopathological abnormalities observed in our DR4tg controls were likely not solely the result of mechanical damage from the i.a. injection itself, since histopathologic scores in PBS-immunized mice receiving PBS i.a were similar to those not receiving the knee injection.

To test whether HomoCitP-induced arthritis was restricted to HLA-DR4, we also examined wild-type B6 mice immunized with HomoCitJED. 50% of B6 mice developed antibody responses against HomoCitJED but none to HomoCitFib and displayed only mild joint histopathological changes. Although some B6 mice mounted detectable immune responses and exhibited mild joint pathology, disease severity was significantly greater in the DR4tg cohort. This disparity suggests that H2-B expressed by antigen presenting cells in wild-type B6 mice have a limited ability to bind and present HomoCitP, whereas DR4tg mice, expressing the SE, present HomoCitP more efficiently, leading to heightened immune activation and resulting in more severe joint damage [4,17,27,28].

In this study, DR4tg mice immunized with HomoCitJED developed immune responses against HomoCitJED and HomoCitFib, consistent with previous findings from our laboratory [4,17]. However, unlike our earlier work where antibody levels declined by day 100, levels in the current study remained stable. This discrepancy may be attributed to the i.a. injections of HomoCitJED, with localized inflammation sustaining autoantibody levels [17]. Our earlier studies also detected epitope spreading, such that antibodies against CitJED were present on days 30, 50, and 70 post-primary immunization, but not day 100. However, in the present study 30% of mice exhibited such responses at timepoints between days 70 and 137 [4,17]. Similarly, in humans, Kolarz *et al.* reported that 47% of AHCPA-positive patients also had ACPAs [29]. The functional significance of this epitope spreading between CitP and HomoCitP in RA progression remains unclear. Mydel *et al.* previously reported that immune responses to CitP alone resulted in only mild disease, whereas responses to HomoCitP were essential for the development of erosive arthritis [26]. We did not observe a correlation between CitJED responses and worse arthritis scores (data not shown), and clinical data similarly indicate that double-positive patients do not necessarily experience greater joint damage [16]. Further investigation into the role of epitope spreading between CitP and HomoCitP in RA disease progression is required.

Although immune responses towards HomoCitJED and HomoCitFib were observed and the antigens are present in the joints of naïve mice, these are insufficient to drive spontaneous arthritis [4,19]. A similar phenomenon is seen in humans, where ACPAs and AHCPAs are detected years before the onset of clinical disease, consistent with the “two-hit” hypothesis of RA [30–32]. According to this theory, the first hit involves the generation of ACPAs and potentially AHCPAs, while the second hit leads to synovial inflammation, which amplifies antigen exposure and leads to joint damage [32]. In our model, systemic immunization with HomoCitJED provides the first hit by priming the immune system. Although we did not detect increased antigen levels in the joint, the knee i.a. injections of HomoCitJED serve as the second hit by inducing local inflammation and transiently elevating local antigen concentrations until clearance occurs, thereby directing the immune response to the joint. This is in part supported by Turunen *et al.* who found that worse disease severity was associated with increased levels of HomoCitP [33].

While these i.a. injections might aid in the triggering of arthritis in our model, we observed that the knee injected with HomoCitJED exhibited joint damage comparable to the contralateral knee injected with PBS. One possible explanation is that antigen clearance by local immune responses leads to systemic responses that drive pathology beyond the injected joint. This phenomenon was previously described in the transforming growth factor β1 injection with treadmill running (TTR) model of joint injury, where i.a. injections of hyaluronic acid did not significantly alter bone structure in the injected knee. Interestingly, structural changes were observed in the contralateral knee instead. Shen *et al.* proposed that this was due to the rapid clearance of hyaluronic acid in the injected knee. [34].

Development of mouse models that better reflect human RA is essential, since current models are frequently critiqued for not reproducing the sex-specific differences seen in patients. In the collagen-induced arthritis (CIA) model, which is one of the most common models used in RA research, male mice experience a greater incidence of arthritis and more severe disease [35,36]. While we found no difference in disease incidence between the male and female mice, there was a non-significant increase in disease severity in males. Potential reasons for this discrepancy could include female mouse sex hormones, since oophorectomy makes female mice as susceptible to CIA as male mice [37]. Additionally, the presence of murine MHC-II chains could play a role, as a different DR4tg mouse strain lacking all endogenous MHC-II chains were found to have greater CIA incidence in females (52%) compared to males (15%) [38].

Other limitations of our study include using only 2D histopathological analysis to assess disease severity. Radiographic methods, such as MRI or CT, would aid in the understanding of changes to the whole knee joint. While our model exhibited the hallmark features of RA joint histopathology in the knee, it did not result in systemic arthritis. Compared to the hind paw of CIA mice, our model had low histologic arthritis severity scores; however, comparison of histopathological damage between RA mouse models is challenging due to differing features and severity [6,39,40]. While our study does have limitations, there are several strengths. We used a synthetic peptide, HomoCitJED, which captures several clinically relevant antigens, such as homocitrullinated fibrinogen. Furthermore, our animal model is physiologically relevant to human RA as it expresses the greatest genetic risk factor for disease development, HLA-DR4. Similarly, the i.a. injections were administered without an adjuvant to more closely mimic the potential pathogenesis of disease.

## 5. Conclusion

In summary, our study supports the ability of homocitrulline-specific immune responses to induce RA-like disease in DR4tg mice following i.a. injections. Additionally, this was observed to be enhanced by expression of HLA-DR4, but not restricted, as some B6 mice developed antibody responses and mild disease. Overall, this data reinforces the pathogenic role of homocitrulline-specific immune responses, providing a potential model for the development of disease-specific therapeutics.

## Acknowledgements

We would like to thank Garth Blacker for his contributions to the animal studies and Patti Kiser for her assistance with the interpretation of histopathological features. We also appreciate the input of Jaspreet Kaur and Sofya Ulanova in reviewing and providing critical feedback on the manuscript.

---

**Supplementary Figure 1.**
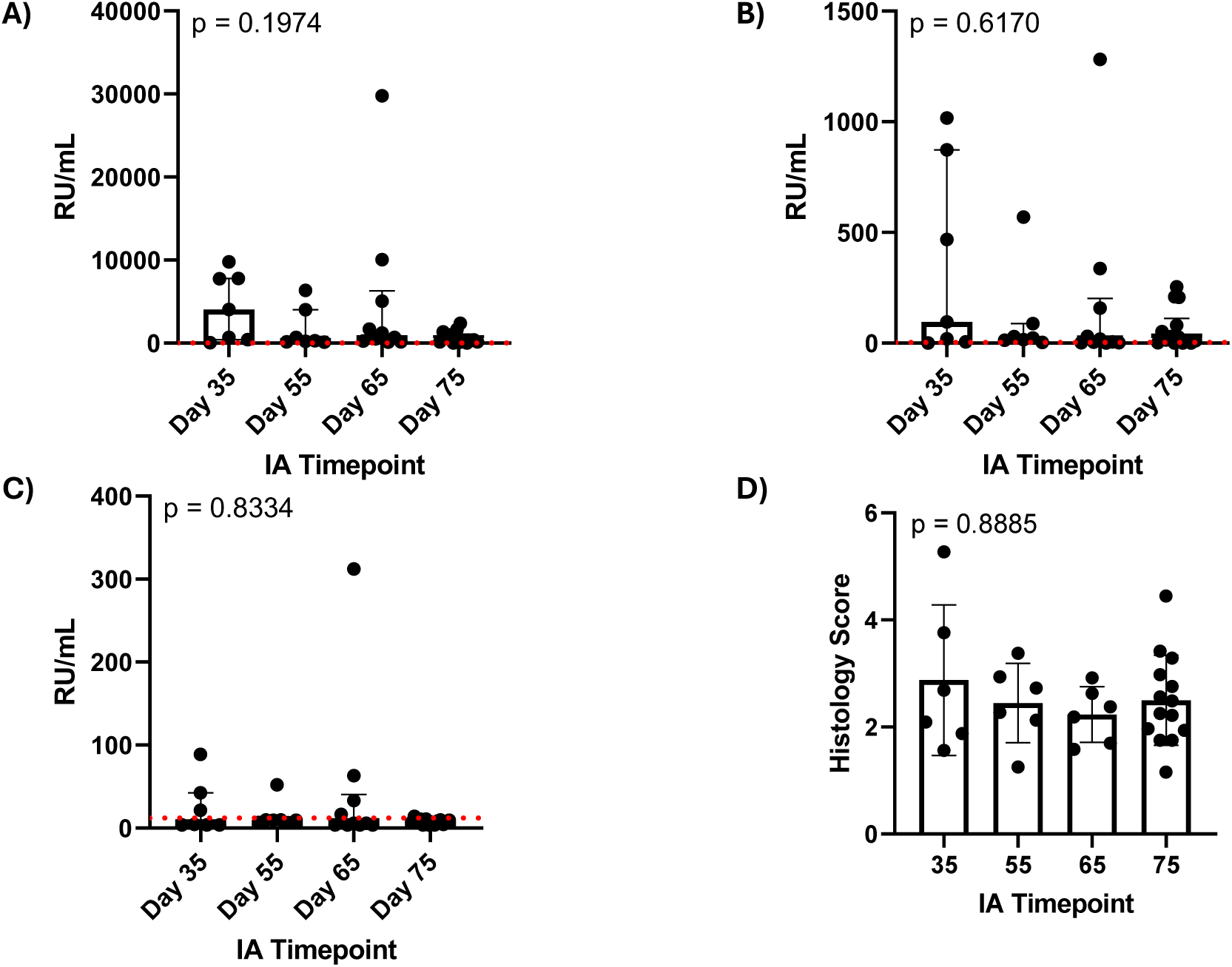
Effect of i.a. timing on antibody responses and arthritis severity. HomoCitJED immunized DR4tg mice were given a total of 3 HomoCitJED i.a. injections, administered at 7-day intervals, with the first injection given on either day 35, 55, 65 or 75. Terminal IgG antibody responses were compared between the i.a. timepoints for (**A**) HomoCitJED, (**B**) HomoCitFib, and (**C**) CitJED. Antibody levels, determined via ELISA, are shown in RU/mL. Graphs display the median (IQR), with each symbol representing one mouse. The positive cutoff is shown by a red dashed line; N = 7-14. Brown-Forsythe ANOVA statistical test of area under the curve was performed. (**D**) Comparison of total histopathological scores between the i.a. timepoints. Histological scores from the right and left knee were averaged. Each symbol represents one mouse with the positive cutoff shown by a red dashed line; N = 6-14. A Kruskal-Wallis statistical test was performed.

**Supplementary Figure 2.**
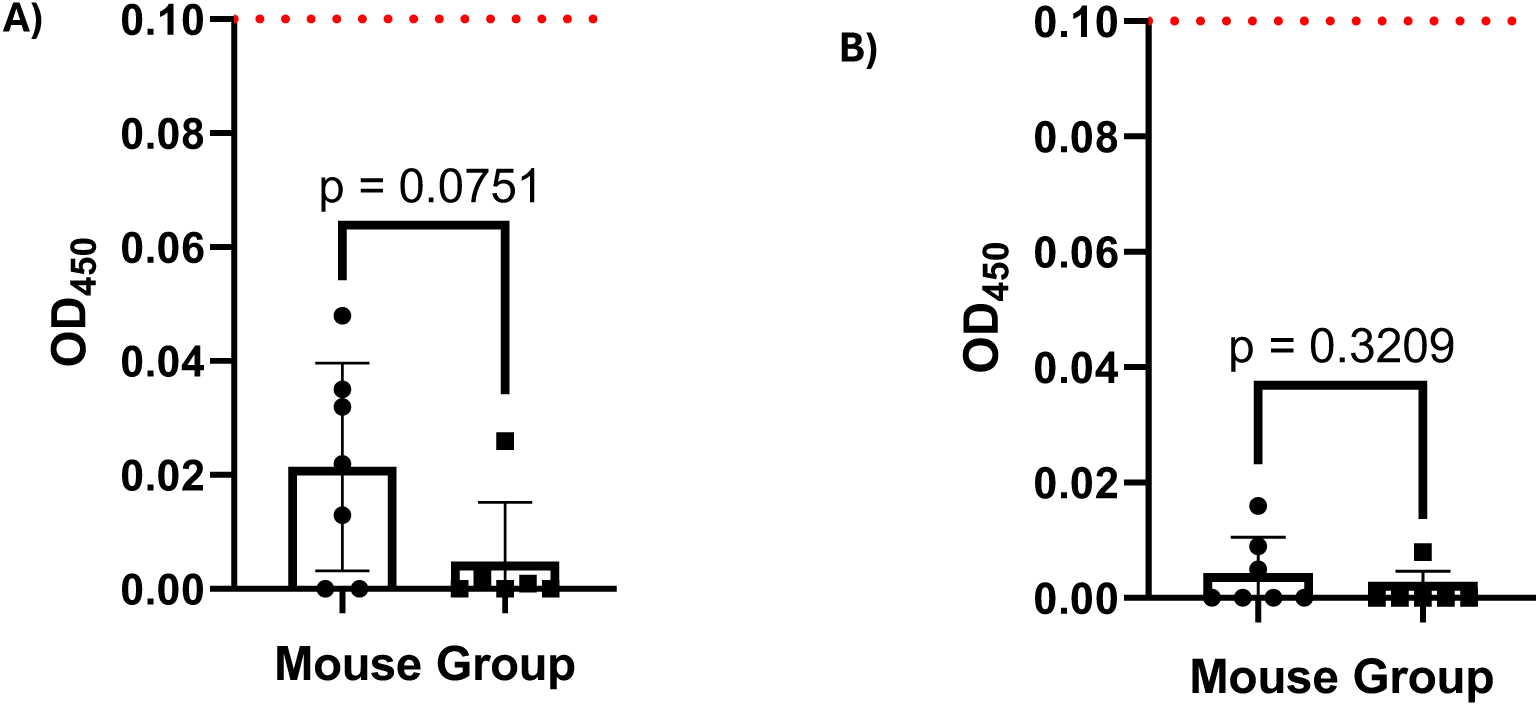
IgG antibody responses from DR4tg and B6 mice. Terminal IgG antibody responses from DR4tg [HomoCitJED immunized (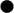)] and B6 [HomoCitJED immunized (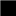)] mice were screened for (**A**) anti-fibrinogen, and (**B**) anti-CitFib antibodies. All mice received i.a. injections starting on day 75 of HomoCitJED in the left knee and PBS in the right knee. Antibody levels, determined via ELISA, are shown in OD at 450 nm (1:100 dilution). Graphs display the mean (SD), with each symbol representing one mouse. The positive cutoff is shown by a red dashed line; N = 6-7. An unpaired t-test was used for anti-fibrinogen and anti-CitFib autoantibodies.

**Supplementary Figure 3.**
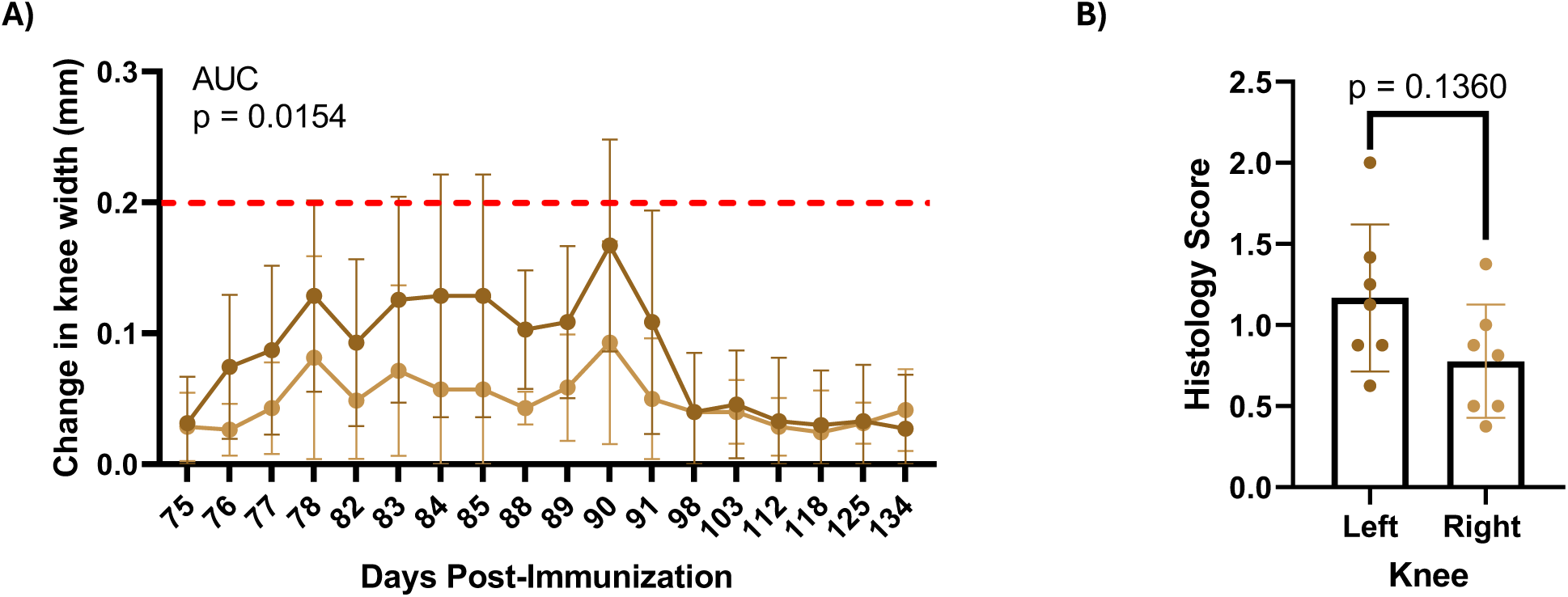
Effect of i.a. injections on swelling and histopathology. (**A**) Change in knee width caliper measurements of PBS immunized DR4tg receiving PBS i.a.’s in the left knee (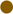) and no i.a. in the right (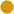). Each symbol represents the mean value (SD) with the swelling cutoff shown by a red dashed line; N = 7. A paired t-test of area under the curve was performed. (**B**) Total histopathological scores were compared between the left (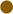) and right (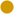) knees of these PBS-immunized DR4tg mice. Each symbol represents one mouse knee; N = 7. A paired t-test was performed.

**Supplementary Figure 4.**
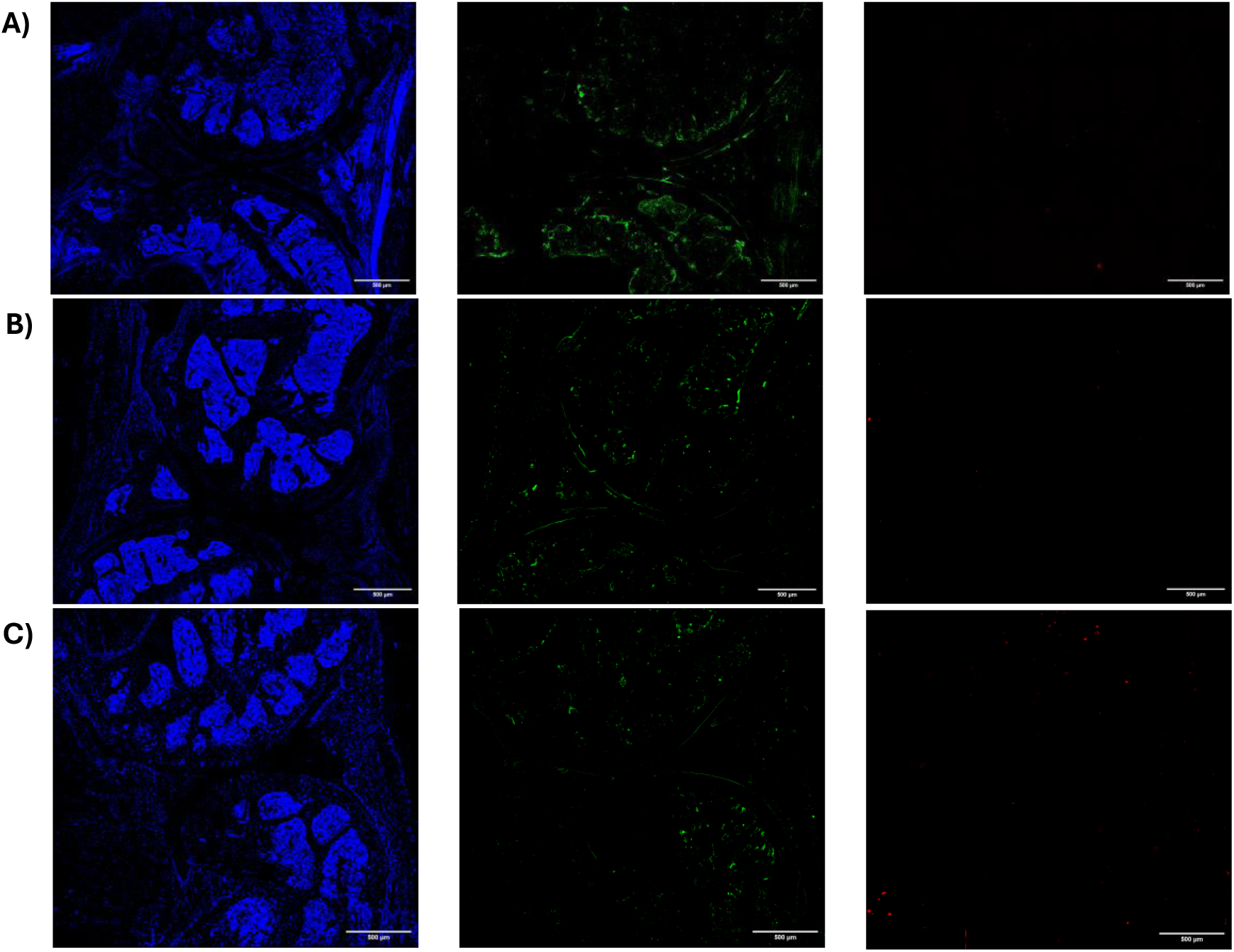
Representative isotype controls of CitP and HCitP in joints of DR4tg mice. Representative isotype images of (**A**) HomoCitJED immunized DR4tg mice, (**B**) PBS immunized DR4tg mice, and (**C**) naïve DR4tg mice. Joints were stained such that nuclei, CitP, and HCitP are DAPI (blue), FITC (green), and CY5 (red), respectively. Immunized mice received i.a. injections of HomoCitJED in the left knee and PBS in the right knee. The images are at 20x magnification; scale bar = 500 µm.

**Supplementary Figure 5.**
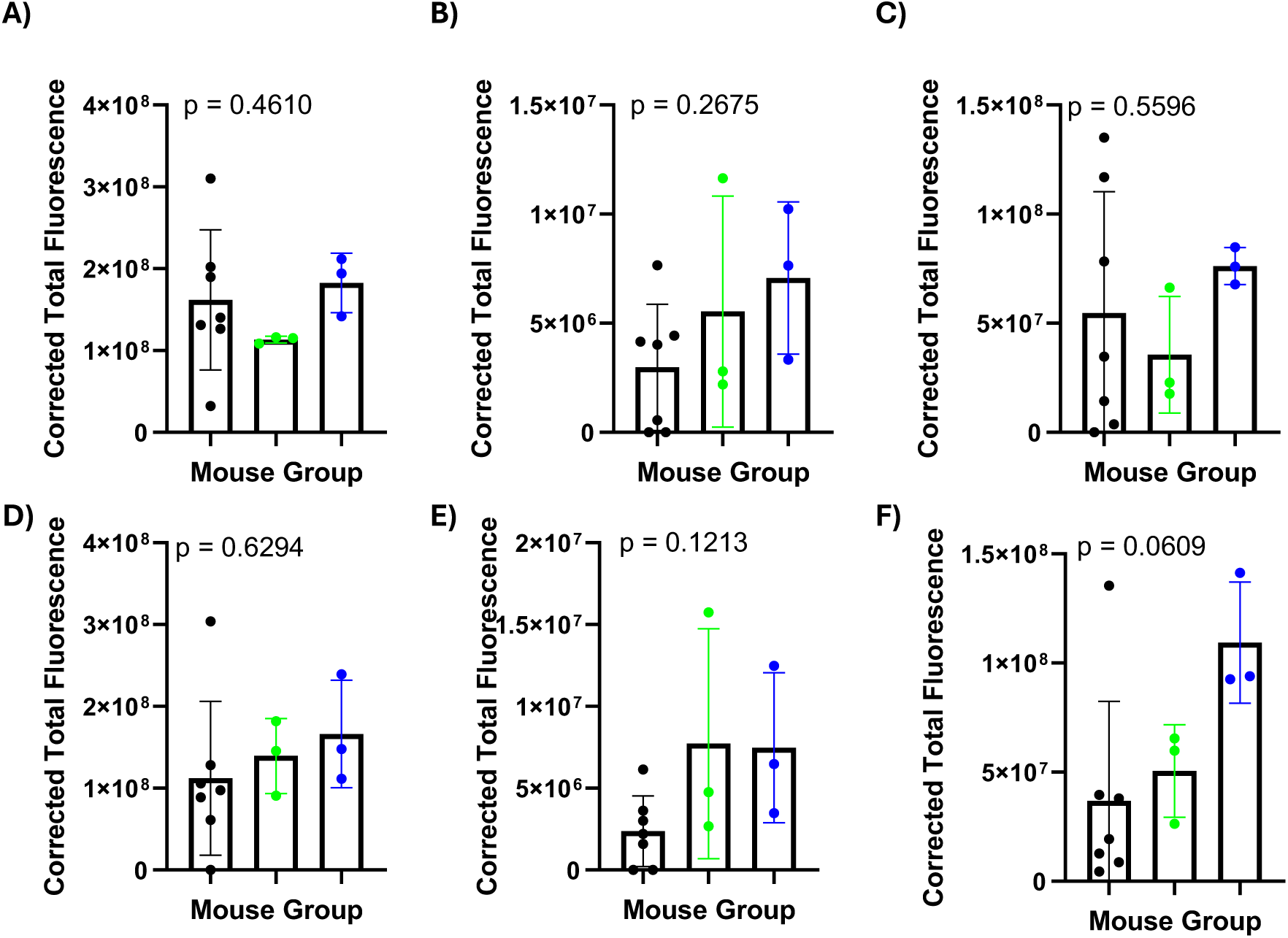
Corrected total fluorescence (CTF) of CitP and HCitP in joints of DR4tg mice. CTF for CitP and HCitP in the bone marrow (**A** and **D**, respectively), cartilage (**B** and **E**, respectively), and synovium (**C** and **F**, respectively) was calculated by subtracting background fluorescence. CitP and HCitP levels in the left knee were compared between HomoCitJED immunized DR4tg mice (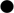), (**B**) PBS immunized DR4tg mice (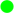), and (**C**) naïve DR4tg mice (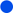). Injected mice received i.a. injections of HomoCitJED in the left knee. Graphs display the mean (SD), with each symbol representing one mouse; N = 3-7. An ordinary one-way ANOVA statistical test was performed.

